# Normative Deviations Reveal Task-Evoked and Clinical Network Reorganization

**DOI:** 10.64898/2026.02.04.703503

**Authors:** Jean-Philippe Kröll, Mohamed Abdelmotaleb, Harun Kocataş, Veronica Müller, Lya Paas, Marcus Meinzer, Agnes Flöel, Simon Eickhoff, Kaustubh R. Patil

**Affiliations:** Institute of Neuroscience and Medicine, Brain & Behaviour (INM-7), Research Centre Jülich, Jülich 52428, Germany; Institute of Systems Neuroscience, Medical Faculty, Heinrich Heine University Düsseldorf, Düsseldorf 40225, Germany; Department of Neurology, University Medicine Greifswald, Greifswald, Germany; German Centre for Neurodegenerative Diseases (DZNE) Standort Greifswald, Greifswald, Germany

## Abstract

Understanding how cognitive demands and pathology reshape large-scale functional connectivity (FC) requires methods that are both multivariate and region-specific. Here we introduce One-class SVM-based Connectome Anomaly Recognition (OSCAR), a normative modelling framework that detects condition-related deviations in the multivariate connectivity profile of a brain region. OSCAR learns the distribution of region-to-whole-brain connectivity patterns given a reference state sample (e.g. resting-state data; RS) using a one-class support vector machine (OCSVM). The trained models are then applied to FC profiles from a target condition (e.g., task or patient group). The outlier proportions are used to quantify the difference between the reference and target condition. We validated OSCAR on three diverse tasks and a patient cohort with early psychosis. OSCAR consistently identified condition-sensitive regions in networks known to support conflict processing, object-location memory, lexical learning, and early psychosis, respectively, including thalamic and basal ganglia regions. Moreover, it detected additional well-established task– or disease-relevant parcels not captured by the comparison method permutation-based multivariate analysis of variance (perMANOVA). Regions flagged by OSCAR were at least as close, and often closer, to independent task-activation findings than those identified by perMANOVA. These results demonstrate that OSCAR provides an interpretable, region-centred normative modelling approach that is sensitive to subtle multivariate FC deviations, and offers a practical tool for mapping condition-specific reconfigurations of functional brain networks with high external validity, in both experimental and clinical settings.

## Introduction

Understanding human brain function requires accounting for two fundamental organizational principles: regional specialization (Kanwisher, 2010) and functional integration (Friston, 2002). Regional specialization describes how distinct brain regions are tailored to support specific functions, such as sensory encoding (Wei et al., 2024), motor planning (Makary et al., 2017), memory formation (D’Esposito et al., 2000), or abstract reasoning (Gilead et al., 2014). Yet these localized functions alone cannot explain human cognition and disease often manifested as coordination across many regions (Bressler and Menon, 2010). Functional integration therefore refers to the context-dependent interactions between several brain regions. Contemporary neuroimaging work has shown that these principles are inherently intertwined: specialized regions contribute to cognition through their embeddedness within large scale networks, while network-level dynamics depend on the differentiated capabilities of individual nodes (Wang et al., 2021).

Crucially, this interplay is not static. The brain continually adapts its functional organization to meet the demands of unfolding cognitive processes, dynamically balancing between segregation and integration (Kolskår et al., 2018; Shine et al., 2016). These dynamics also imply that the functional organization can be (systematically) perturbed. Tasks or external inputs (such as naturalistic stimuli) place distinct demands on regional processing, and any local change is expected to drive corresponding changes in large-scale connectivity. Conversely, altered connectivity necessarily modifies the functional role of individual regions. Understanding how brain organization is modulated by a given context is critical for elucidating the neural basis of behavior as well as clinical conditions. The present work proposes a new method to detect changes in brain functional organization based on how each region is typically embedded within whole brain networks.

While both activation-based and conventional univariate functional connectivity (FC) analyses can reveal condition-specific effects, each captures only part of the underlying reconfiguration (Murphy et al., 2016; Satake et al., 2024). Activation studies focus on local signal changes but do not describe how these local effects alter a region’s embedding within large-scale networks. Univariate FC analyses, in turn, capture changes in individual connections but do not consider multivariate differences. Methods to identify multivariate FC differences have been proposed (Etzel et al., 2013; Kriegeskorte, 2008; Nieto-Castanon, 2022). Those methods, however, compare conditions or groups, without providing a reference model of what a region’s connectivity profile is expected to look like under a normative condition. On the other hand, data-driven approaches such as independent component analysis (ICA) provide information about large-scale functional networks, but they do not directly test whether specific regions alter their connectivity patterns in a condition-dependent manner (Huang et al., 2024; Mckeown et al., 1998). Multivariate normative modelling complements these approaches by providing a reference expectation for each region’s typical connectivity profile, enabling deviations from this normative embedding to be quantified in a spatially specific and context-sensitive manner (Marquand et al., 2016; Rutherford et al., 2023). As a result, these frameworks have the potential to uncover reorganizations that may be subtle, distributed, or not well captured by other methods. Thereby, these approaches can help characterize how tasks reorganize network structure and how external perturbations, such as non-invasive brain stimulation or pharmacological interventions, modify these networks, even under heterogeneous or less controlled conditions. For example, in clinical populations, disease processes often alter the brain in diffuse or individually variable ways that need not align with group-level contrasts (Segal et al., 2023). Deviations from normative embedding might identify atypical regional roles or early, preclinical disconnection patterns that are not detected by conventional approaches. In addition, a normative model can be used to obtain individual-level information on new individuals. Deviations from normative network embeddings thus provides a principled means of identifying where and how functional organization shifts under different cognitive or sensory demands.

We here propose a new method, One-class SVM-based Connectome Anomaly Recognition (OSCAR), that uses a sophisticated multivariate density estimation method (OCSVM), and considers region-to-whole brain patterns of connectivity. It provides a p-value for each region that can be used to identify regions whose multivariate connectivity pattern differs in the target condition compared to the reference. We here consider the identification of regions affected by a given target condition an outlier detection problem, treating healthy resting-state (RS) as the reference state and aiming to detect outliers within task or patient data. To validate OSCAR, we applied it to the data from the Amsterdam Open MRI Collection (AOMIC, resting state and gender stroop task) (Snoek et al., 2021) and two task datasets from the *Modulation of brain networks for learning and memory by transcranial electrical brain stimulation: A systematic, lifespan approach* (MeMoSLAP) (https://www.memoslap.de/de/forschung/). Here, we expected OSCAR to identify task-relevant regions, based on the assumption that regions engaged by a given task should exhibit changes in their connectivity profiles. Further, we tested OSCAR on data from patients with early psychosis from the Human Connectome Project for Early Psychosis (HCP-EP) (Jacobs et al., 2024). Based on previous findings that psychosis is accompanied by a disruption in RS functional connectivity (rsFC) (Cattarinussi et al., 2024; Dong et al., 2018), we anticipated that OSCAR would detect psychosis-specific FC abnormalities that diverge from the normative FC patterns observed in healthy individuals. We compared our results to a standard Multivariate Analysis of Variance based on permutation testing (perMANOVA, (Anderson, 2001).

## 2. Methods

### 2.1 Datasets

fMRI data from two open datasets were used. The first dataset, AOMIC, consists of three sub datasets—PIOP1, PIOP2, and ID1000—containing multimodal 3T MRI data from healthy individuals, including structural and functional MRI data. Data from PIOP1 (N=226, 129 females, mean age 22.2 ± 1.8 years) and PIOP2 (N=216, 120 females, mean age 21.9 ± 1.79 years), were used. The RS data from PIOP2 was used as the reference while the Gender-Stroop task (GStroop) from PIOP1 was used as the target condition. The gender-Stroop task probed cognitive conflict and control processes. Participants viewed images of twelve male and twelve female faces paired with either congruent or incongruent gender labels (Dutch words for “man,” “sir,” “woman,” and “lady,” shown in upper or lower case) positioned above each face. Trials followed a randomized event-related design. The task lasted 490 seconds (245 volumes, TR = 2 s).

In addition, RS and task fMRI data of the preparatory phase of a larger, multicenter, double-blinded, sham-controlled crossover tDCS study, *Modulation of brain networks for memory and learning* (MeMoSLAP), was used (https://www.memoslap.de/de/forschung/), restricted to imaging sessions where sham stimulation was applied. Data for two different tasks, object-location-memory task (OLM, N=20, 10 females; mean age 25 +/− 5.56, (Abdelmotaleb et al., 2025)) and novel-word-learning task (NWL, N=20, 10 females; mean age = 26 +/− 5.86 (Kocataş et al., 2025)), were taken from two different RU projects, scanned at the same scanner using the same acquisition parameters. For training, RS data from the AOMIC dataset from PIOP1 and PIOP2 were combined for training (N = 442), and task differences were assessed separately for each target condition (OLM and NWL). The OLM task involves learning associations between objects (houses on a two-dimensional [2D] map) and their locations through repeated presentations, utilizing both instruction– and feedback-based mechanisms, using a block design with four learning stages (for more details refer to (Abdelmotaleb et al., 2025)). For the OLM analysis, only data during the six task blocks was used for further analysis. The NWL task required learning picture–pseudoword associations across six blocks, each comprising four learning stages. In each learning block, participants were required to identify correct combinations of five object pictures and pseudowords across four learning stages. Pictures were presented twice in each learning stage (10 trials) and were paired with either a correct or incorrect pseudoword (for more details refer to (Kocataş et al., 2025)). For the analysis of the NWL task, only data during the six task blocks was used in the further analysis.

We also used data from the Human Connectome Project for Early Psychosis (HCP EP), including RS data from patients with early psychosis (N=184, 70 female, mean age 23.1 +/−3.7) and healthy controls (N=68, 24 female, mean age 24.4 +/− 4.3). Here, training relied on RS data from the healthy controls, while patients’ data was used as target condition.

In all cases, the same data splits were used to detect differences by perMANOVA.

### 2.2 Data Preprocessing

For AOMIC, preprocessed data using fMRIPrep version 1.4.1 (Esteban et al., 2019) provided via OpenNeuro was used. In brief, data were motion corrected using mcflirt (FSLv5.0.9, (Jenkinson et al., 2002)) followed by distortion correction by co-registering the functional image to the respective T1 weighted image with inverted intensity (Wang et al., 2017) with six degrees of freedom, using bbregister (FreeSurfer v6.0.1). In a following step, motion correction transformations, field distortion correction warp, BOLD-to-T1-weighted transformation and the warp from T1-weighted to MNI were concatenated and applied using antsApplyTransforms (Avants et al., 2011) using Lanczos interpolation (Snoek et al., 2021). Mean white-matter, CSF, and global signals (including their derivatives and squared terms under the full strategy) were regressed out to remove non-neuronal and global physiological noise.

Data from MeMoSLAP was preprocessed with fMRIPrep 24.1.1 (Esteban et al., 2019). In brief, each BOLD run was corrected for head motion using MCFLIRT (FSL, (Jenkinson et al., 2002)). The BOLD reference was co-registered to the corresponding T1-weighted structural image using FreeSurfer’s mri_coreg, refined with FLIRT boundary-based registration (six degrees of freedom (Greve and Fischl, 2009)). Spatial normalization to MNI152NLin2009cAsym was performed via nonlinear registration using ANTs (Greve and Fischl, 2009). All spatial transforms—including motion correction, susceptibility distortion correction (when available), BOLD-to-T1w alignment, and T1w-to-MNI warps—were combined and applied in a single interpolation step using cubic B-splines. Mean white-matter, CSF, and global signals (including their derivatives and squared terms under the full strategy) were regressed out to remove non-neuronal and global physiological noise.

The HCP EP dataset was preprocessed with the minimal preprocessing HCP pipeline (Glasser et al., 2013). The HCP minimal functional preprocessing pipelines consist of correction of gradient-nonlinearity-induced distortion, realignment of the time-series to correct for subject head motion, registration of the fMRI data to the structural data, reduction of the bias field, normalization of the 4D image to a global mean, masking the data with the final brain mask, and the spatial smoothing using a novel geodesic Gaussian surface smoothing algorithm with 2 mm FWHM. Following minimal preprocessing, ICA-FIX (Salimi-Khorshidi et al., 2014) was applied to identify and remove noise components using an automated classifier. Mean white-matter, CSF, and global signals (including their derivatives and squared terms under the full strategy) were regressed out to remove non-neuronal and global physiological noise.

### 2.4 OSCAR

To create region-specific normative models, we here propose a new method, One-class SVM-based Connectome Anomaly Recognition (OSCAR), that uses a multivariate density estimation method (OCSVM), and considers region-to-whole brain connectivity patterns. Consider the case where we want to identify differing connectivity patterns between two Groups, A (reference state, e.g. RS) and B (modulated state, e.g. task or patient data). For a region we train a OCSVM (Schölkopf et al., 1999) classifier using the group A data to learn the multivariate density describing the connectivity pattern of that region with all other regions. We then apply this OCSVM model to the connectivity pattern of the same region for individuals in both group A and group B. This provides the inliers (individuals whose connectivity pattern conforms with the OCSVM density estimation) and outliers (individuals whose connectivity pattern diverges) for the two groups. A chi^2^ test is then applied to compare the inliner and outlier proportion of the two groups. Group B will have a higher proportion of outliers if it has a different multivariate profile than group A resulting in a low p-value. By repeating this procedure for every region and controlling for multiple testing using Bonferroni correction (Dunn, 1961), we identify regions where the two groups differ.

In this study, we considered the identification of regions affected by a given target condition an outlier detection problem, treating healthy RS as the reference state and aiming to detect outliers within task or patient data.To prepare the input features, we first computed FC matrices using regions defined by the 200-resolution Schaefer parcellation (Schaefer et al., 2018) and the 32 subcortical Melbourne parcellation (Tian et al., 2020). Each resulting 232 × 232 Pearson’s correlation matrix were Fisher r-to-z transformed (Fisher, 1915) and then decomposed into 232 region-specific “connectivity profiles.” The connections representing diagonal values in the FC matrix were removed as they show no variance across subjects. Connectivity profiles from the reference state served as input data for training the OCSVM models. This allowed each OCSVM to model the multivariate density underlying that region’s connectivity with the rest of the brain (in the reference state). We utilized sklearn’s OneClassSVM (Schölkopf et al., 1999) with a linear kernel, due to its ability to perform outlier detection in a high dimensional space. The hyperparameter nu (ν) of the OCSVM was optimized using a 5-fold GridSearchCV, searching over the range [0.001, 0.01, 0.025, 0.05, 0.1, 0.25, 0.5]. Νu defines the maximum fraction of training samples allowed to lie outside the decision boundary and the minimum fraction of support vectors that define it. A custom scoring function was used to maximize the number of inliers in the training data. The trained OCSVM models were then applied to connectivity profiles of the target condition individuals. Following the statistical analysis and multiple testing correction, this resulted in a list of regions where the target condition is statistically different from the reference.

### 2.4 Comparison with perMANOVA

We compared OSCAR with perMANOVA based on Anderson (Anderson, 2001). We selected perMANOVA for comparison as it is designed for detecting multivariate group differences. perMANOVA performs a distance-based group comparison that tests whether the average multivariate distances between conditions exceed within-condition variability using permutation testing. The following procedure was applied to each region’s connectivity profile. The connections representing diagonal values in the FC matrix were removed as they show no variance across subjects and remaining connections were Fisher r-to-z transformed (Fisher, 1915). A Euclidean distance matrix was then computed between all pairs of samples, i.e. the FC connectivity profiles of all subjects from two given conditions. Next, a null distribution is generated by permuting the sample assignment to the two conditions multiple times, and the distances were recalculated. A pseudo-F statistic is then calculated by comparing the observed between-group distances to the within-group distances. Finally, the p-value is determined by comparing the observed test statistic to the permutation-derived null distribution. We conducted 5000 permutations in this analysis, repeated for each region and controlling for multiple comparisons using Bonferroni correction. In addition, we compared OSCAR to an adapted version of functional connectivity MultiVariate Pattern Analysis (fc-MVPA), a method for detection of multivariate patterns in FC described by Nieto-Castanon (Nieto-Castanon, 2022). However, in the datasets used here, fcMVPA did not identify any significant differences. This may be due to our region-wise adaptation of the method, which was originally designed for voxel-wise analyses.

### 2.6 Validation Against Prior Task-Specific Activation Findings

As there is no single, commonly established method for validating multivariate differences in FC, we relied on two evaluations. First, we quantified the spatial correspondence between the regions detected by a multivariate method (OSCAR or perMANOVA) and regions involved in a respective target condition based on respective task activation analysis or meta analysis. To this end, we used the Euclidean distance of the centroid of the identified regions to the closest (meta-analytic) activation peak as an interpretable similarity metric. This approach provides a reference for assessing whether identified regions lie close to brain areas reliably implicated in the task or, conversely, fall farther away from those regions. For each significant region, only the distance to the closest peak coordinate was considered to quantify its spatial correspondence. Similar spatial correspondence was also used to compare the results of OSCAR and perMANOVA. For the GStroop task, both contrasts of the whole-brain z-value map from the technical validation section of the AOMIC paper were used (Snoek et al., 2021). For the OLM task, we used peak coordinates reported by Abdelmotaleb et al. (Abdelmotaleb et al., 2025) from significant clusters identified in a whole-brain fMRI analysis conducted on the same subjects included in the present study. The spatial correspondence to both contrasts (learning > control and control > learning) was assessed. For the NWL-task, the peak coordinates from a meta-analysis on lexical learning was used for comparison (Tagarelli et al., 2019). When OSCAR and perMANOVA differed in how many regions they detected, we downsampled the method with more findings to its most significant regions to ensure a fair, equal-sized comparison against task activation.

## 3. Results

To provide anatomical interpretations of Schaefer (and Tian) parcels, we mapped each parcel to the Harvard-Oxford (HO) cortical and subcortical atlases (Desikan et al., 2006) by computing voxelwise spatial overlap in MNI space. For each parcel, the HO region with the highest overlap fraction was assigned as its anatomical match. In the following we report Schaefer/Tian parcels as well as the corresponding HO label.

### 3.1 RS vs GStroop Task

For the GStroop task, out of 232 regions OSCAR identified 110 regions while perMANOVA found 56 regions to be significantly different (Fig. 1). OSCAR and perMANOVA shared 32 regions (Fig. 2a). The right-hemisphere precuneus (Schaefer: RH_ContC_pCun_2, HO: Precuneous Cortex), the right anterior globus pallidus (Tian: aGP-rhm, HO: Right Pallidum), and part of the left-hemisphere somatomotor A network (Schaefer: LH_SomMotA_3, HO: Postcentral Gyrus) were the three most significantly different regions found by OSCAR (Supplementary Table 1). The perMANOVA identified a parcel of the right-hemisphere dorsal attention network’s frontal eye fields and two parcels in the ventral prefrontal cortex of the left-hemisphere, belonging to the default network (Supplementary Table 2). Spatial correspondence between the findings of both methods was on average low, with nearest-neighbor distances ranging from 0 to 50 mm (mean ≈ 15 mm) (Fig. 2b). For the negative GStroop contrast (Fig. 2c), the spatial correspondence between regions identified by the two methods and the nearest activation peak did not differ significantly. For the positive GStroop contrast (Fig. 2d), OSCAR-identified regions were significantly closer to the respective activation peaks, indicating a stronger spatial correspondence relative to perMANOVA (Mann-Whitney U test, p = 0.00349).

**Fig. 1.**
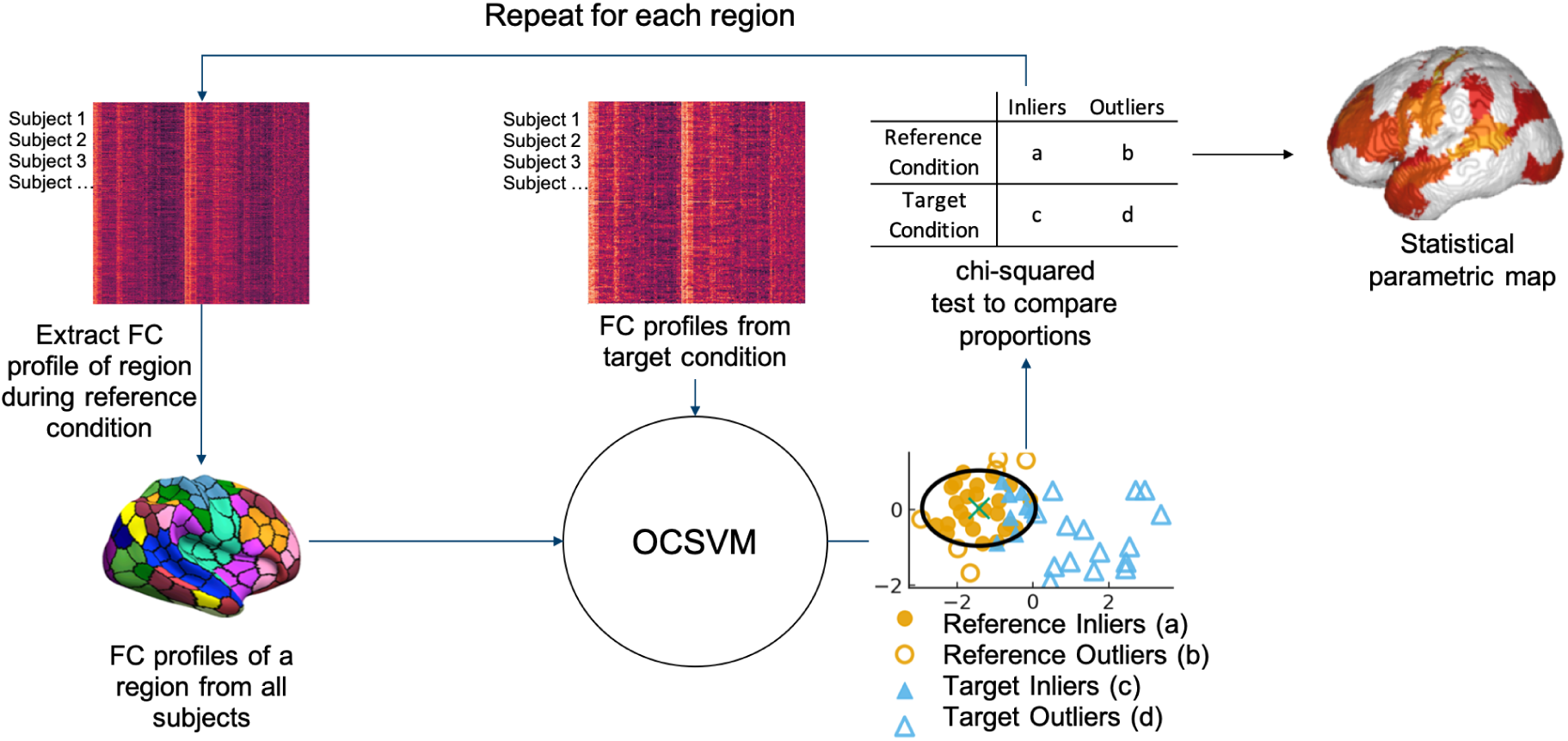
Schematic representation of OSCAR. In the first step, the connectivity matrix is calculated for each subject of the RS data, separately. Then, the connectivity profile of a region (i.e. the connectivity of that region with every other region) is taken from all subjects and used to train a OCSVM-Classifier. This process is repeated for every region, resulting in one model per region. In the second step, the OCSVM-models are applied to connectivity profiles from a target condition (e.g. task– or patient data). Then, the differences between the two groups (RS vs target condition) is statistically quantified and projected onto a brain map. (OSCAR: One-class SVM-based Connectome Anomaly Recognition, OCSVM: One-Class Support Vector Machine, FC: Functional Connectivity)

**Fig. 1.**
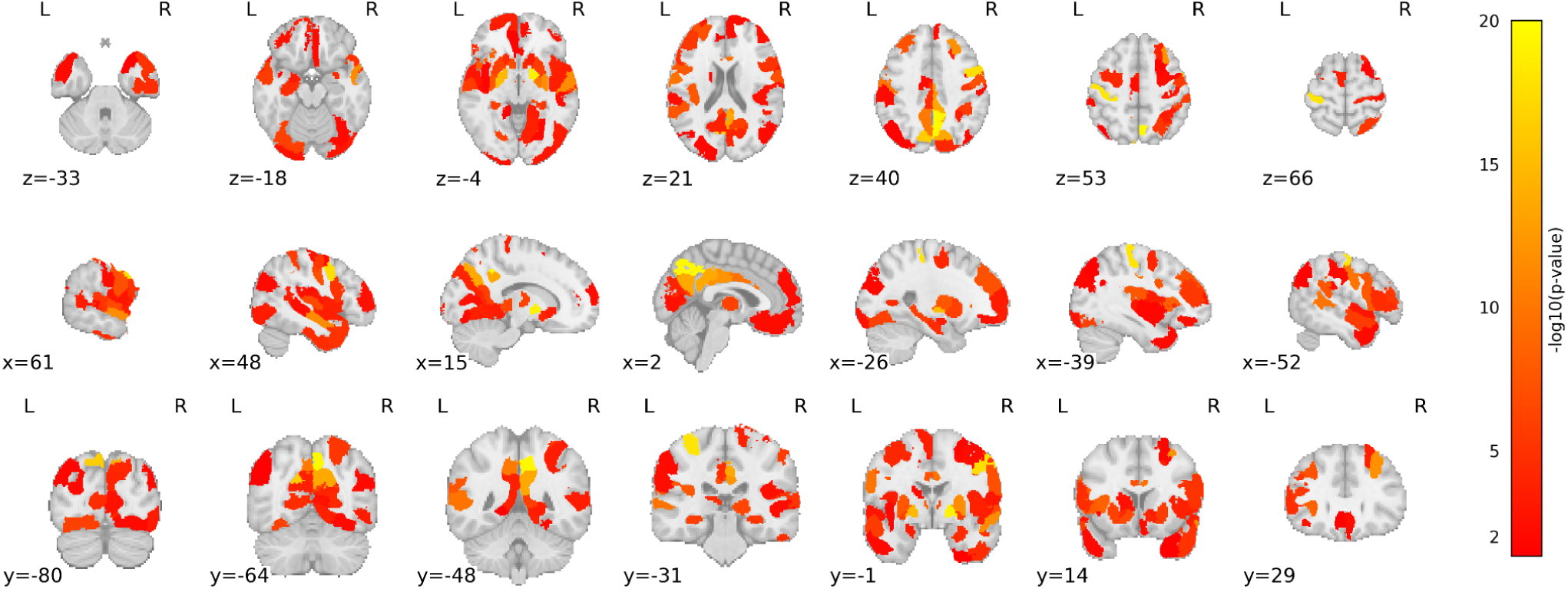
Brain regions with significantly different connectivity patterns between RS and the GStroop task, as identified by OSCAR (L = left, R = right, x = sagittal, y = coronal, z = axial).

**Fig. 2.**
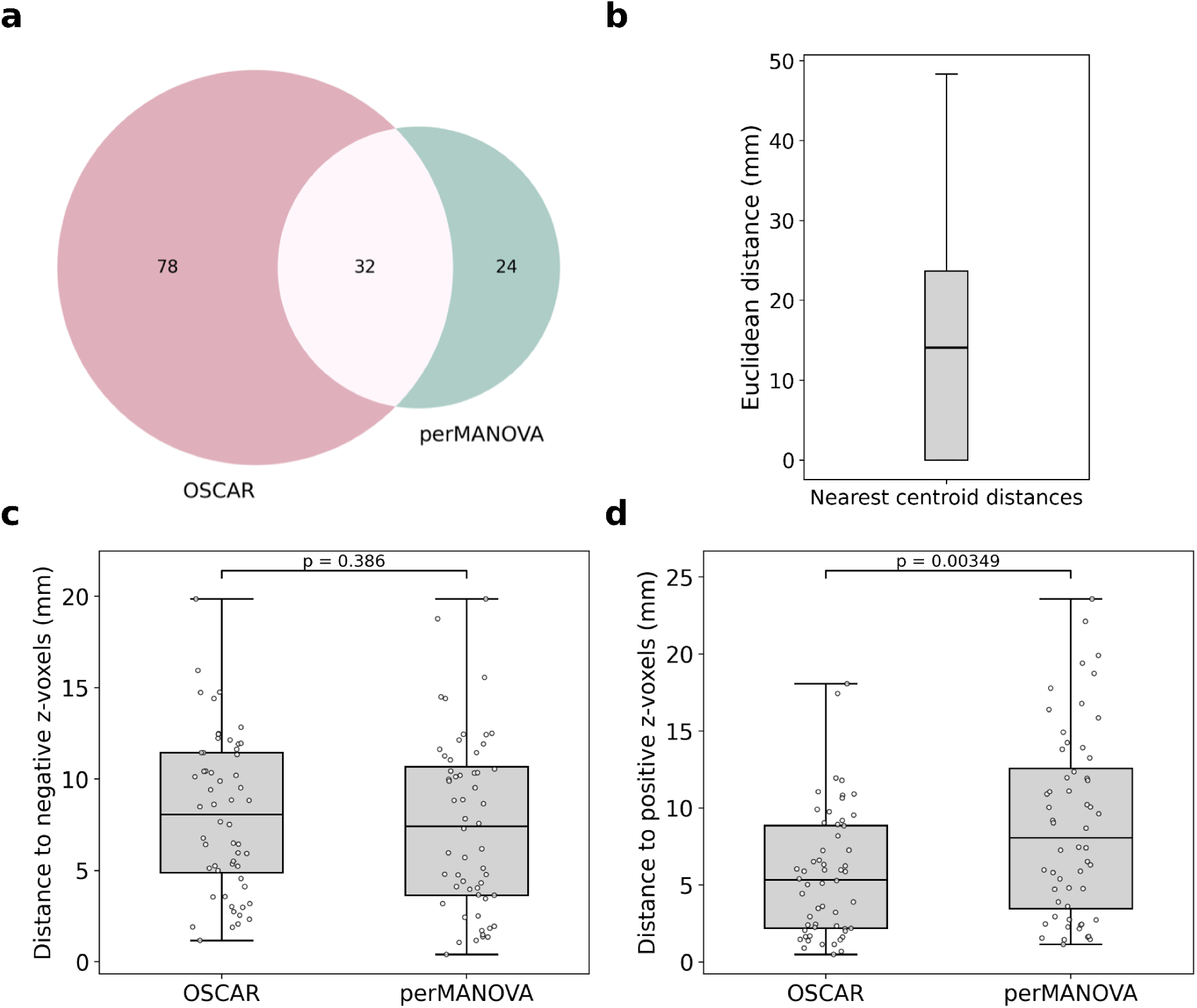
a) Overlap between significant regions identified by OSCAR and perMANOVA. The left circle (pink) represents the 78 regions detected only by OSCAR, the right circle (green) represents the 24 regions only identified by perMANOVA, and the intersection shows the 32 regions jointly identified by both methods. b) Spatial correspondence between the two methods. The plot displays the minimum Euclidean distance from each region identified by one method to the nearest region identified by the other. c) Euclidean distance from each region’s centroid to the nearest activation peak in the negative GStroop contrast (correct > incorrect), with shorter distances indicating greater spatial correspondence. d) Euclidean distance from each region’s centroid to the nearest activation peak in the positive GStroop contrast (incorrect > correct), with shorter distances indicating greater spatial correspondence.

### 3.2 RS vs OLM Task

OSCAR identified 48 regions with deviating connectivity in the OLM task compared to RS (Fig. 3). The perMANOVA found 37 significantly differing regions. OSCAR and perMANOVA shared 18 regions (Fig. 4a). The right-hemisphere visual peripheral network’s extrastriate inferior (Schaefer: RH_VisPeri_ExStrInf_1, HO: Lingual Gyrus), striate calcarine (Schaefer: RH_VisPeri_StriCal_1, HO: Intracalcarine Cortex), and occipital pole (Schaefer: RH_VisCent_ExStr_5, HO: Occipital Pole) regions showed the most pronounced state-dependent changes according to OSCAR (Supplementary Table 3).

**Fig. 3.**
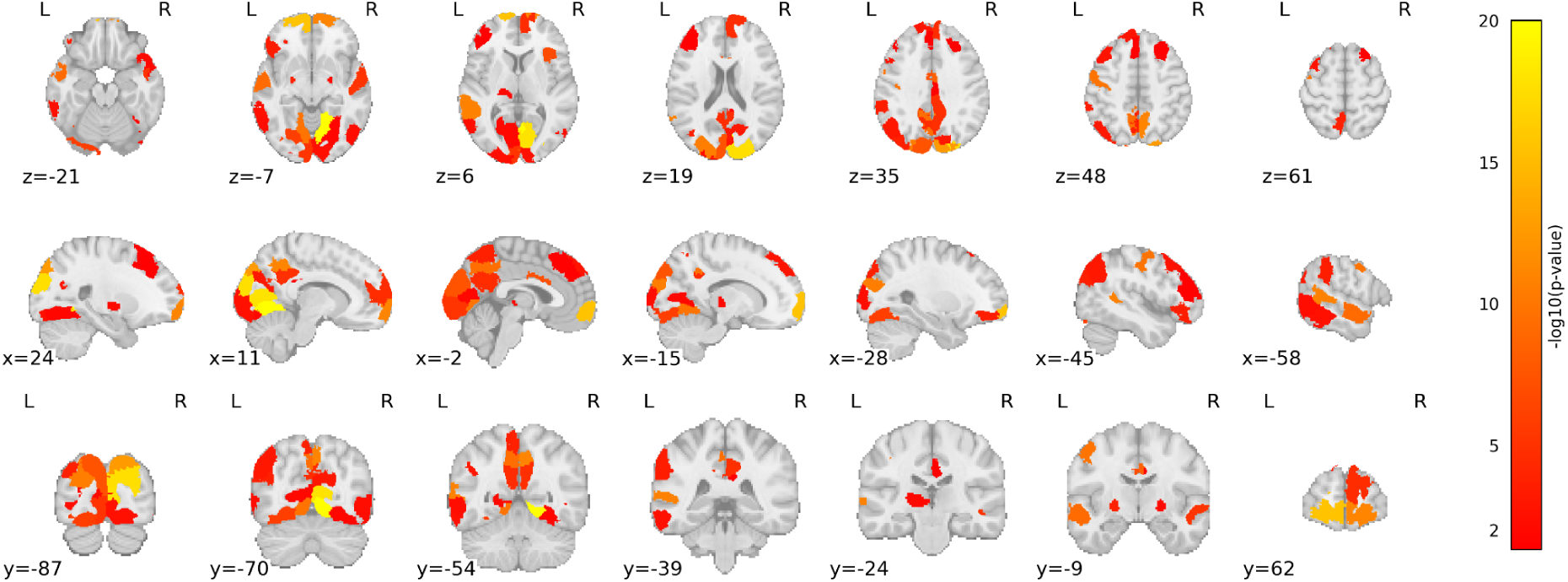
Brain regions with significantly different connectivity patterns between RS and the OLM-task, as identified by OSCAR (L = left, R = right, x = sagittal, y = coronal, z = axial).

**Fig. 4.**
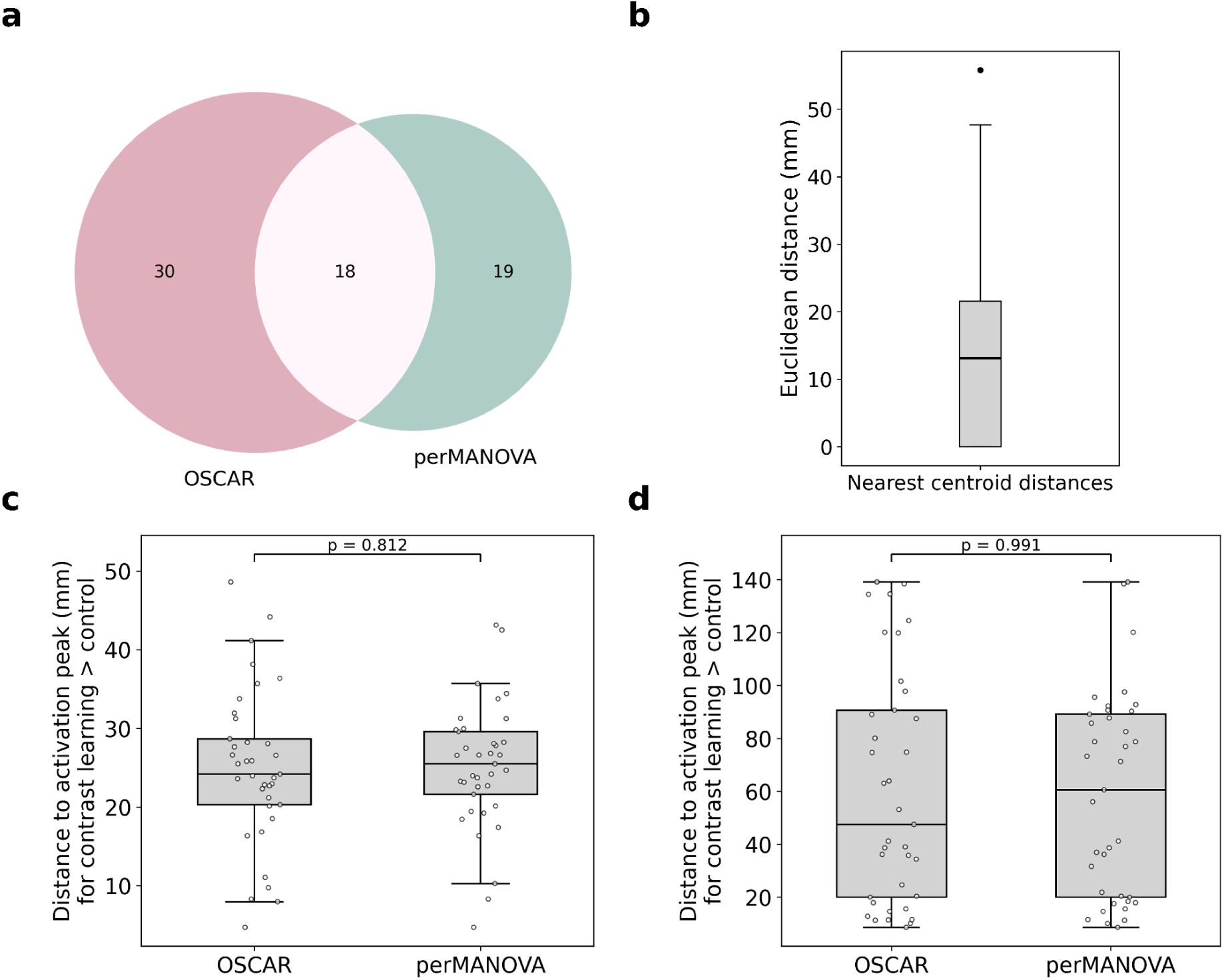
a) Overlap between significant regions identified by OSCAR and perMANOVA. The left circle (pink) represents the 30 regions detected only by OSCAR, the right circle (green) represents the 19 regions only identified by perMANOVA, and the intersection shows the 18 regions jointly identified by both methods. b) Spatial correspondence between the two methods. The plot displays the minimum Euclidean distance from each region identified by one method to the nearest region identified by the other. c) Euclidean distance from each region’s centroid to the nearest activation peak in the OLM task for the contrast learning > control, with shorter distances indicating greater spatial correspondence. d) Euclidean distance from each region’s centroid to the nearest activation peak in the OLM task for the contrast control > learning, with shorter distances indicating greater spatial correspondence. Note that the control > learning contrast only includes five regions.

The perMANOVA identified the left planum porale (Schaefer: LH_SomMotB_Aud_1, HO: Planum Porale), the cingulate gyrus (Schaefer: RH_SalVentAttnA_ParMed_1, HO: Cingulate Gyrus, posterior division) and the central opercular cortex (Schaefer: LH_SomMotB_S2_2, HO: Central Opercular Cortex) as the three most significant regions (Supplementary Table 4). Spatial correspondence between the findings of both methods was moderate, with nearest-neighbor distances ranging from 0 to >50 mm (mean ≈ 15 mm) (Fig. 4b). For both contrasts (learning > control and control > learning), OSCAR-identified regions were on average closer to the respective activation peaks, although not reaching significance (Fig. 4c and 4d).

### 3.3 RS vs NWL-task

For the NWL task, OSCAR identified 18 regions with deviating connectivity, perMANOVA found 17 significantly different regions (Fig. 5). OSCAR and perMANOVA shared three regions (Fig. 6). OSCAR showed highest significance in the left-hemisphere superior temporal gyrus (Schaefer: LH_DefaultB_Temp_3, HO: superior temporal gyrus, anterior division), the left globus pallidum (Schaefer: pGP-lh, HO: left pallidum) and the lateral occipital cortex (Schaefer: LH_DefaultC_IPL_1, HO: lateral occipital cortex, superior division, (Supplementary Table 5). The perMANOVA identified the superior temporal gyrus (Schaefer: LH_DefaultB_Temp_3, HO: superior temporal gyrus, anterior division), the precuneous cortex (Schaefer: RH_DorsAttnB_PostC_3, HO: Precuneous Cortex) and lateral occipital cortex (Schaefer: RH_VisPeri_ExStrSup_3, HO: lateral occipital cortex, superior division) as the most significant regions (Supplementary Table 6). Spatial correspondence between the findings of both methods was low, with nearest-neighbor distances ranging from 0 to >50 mm (mean ≈ 20 mm) (Fig. 6b). Both methods exhibited similarly low proximity to the activation peaks of the meta-analysis (Fig. 6c).

**Fig. 5.**
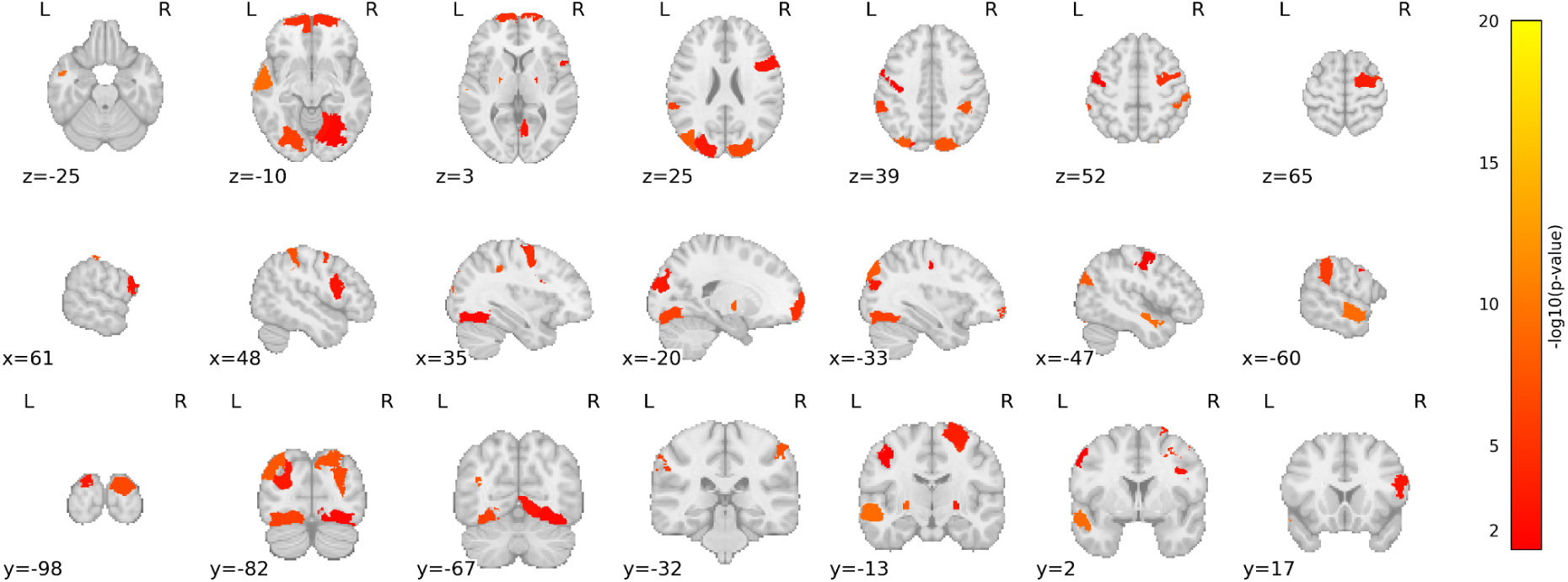
Brain regions with significantly different connectivity patterns between RS and the NWL-task, as identified by OSCAR (L = left, R = right, x = sagittal, y = coronal, z = axial).

**Fig. 6.**
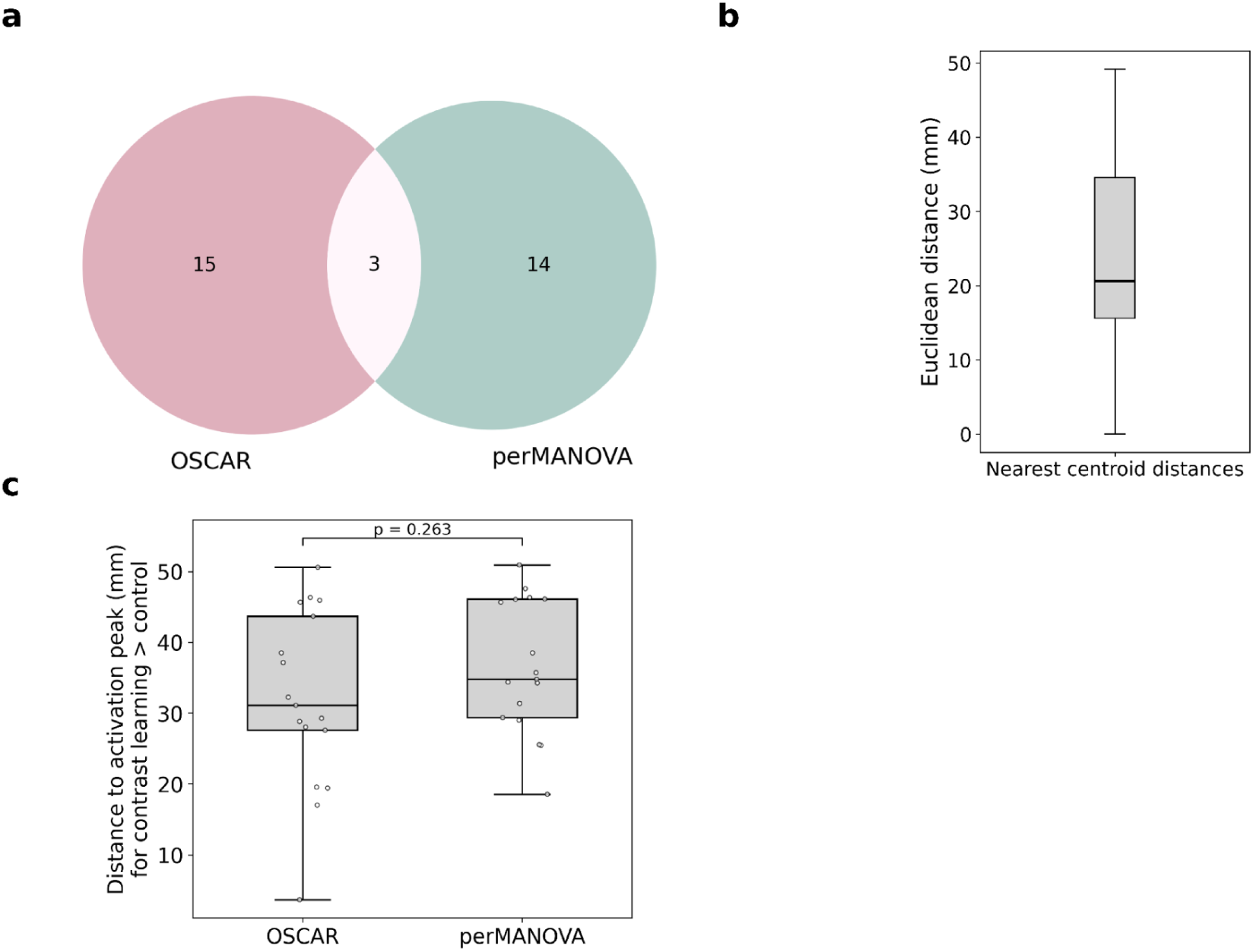
a) Overlap between significant regions identified by OSCAR and perMANOVA. The left circle (pink) represents the 15 regions detected only by OSCAR, the right circle (green) represents the 14 regions only identified by perMANOVA, and the intersection shows the 3 regions jointly identified by both methods. b) Spatial correspondence between the two methods. The plot displays the minimum Euclidean distance from each region identified by one method to the nearest region identified by the other. c) Euclidean distance from each region’s centroid to the nearest peak coordinates from the meta-analysis on lexical learning, with shorter distances indicating greater spatial correspondence.

### 3.4 RS of healthy controls vs RS of EP patients

On data from patients with EP, OSCAR identified 10 regions with deviating RS connectivity compared to RS from healthy controls (Fig. 7). The perMANOVA found 3 significantly different regions, all of which were also identified by the OSCAR. OSCAR showed highest significance in the right-hemisphere ventral anterior thalamus (Tian: THA-VA-rh, HO: Right Thalamus), the left-hemisphere dorsal anterior thalamus (Tian: THA-DA-lh, HO: Left Thalamus), and the right posterior globus pallidus (Tian: pGP-rh, HO: Right Pallidum) (Supplementary Table 7). The three regions identified by the perMANOVA were the left-hemisphere dorsal anterior thalamus, the right-hemisphere dorsal anterior thalamus, and the left-hemisphere ventral anterior thalamus (Supplementary Table 8).

**Fig. 7.**
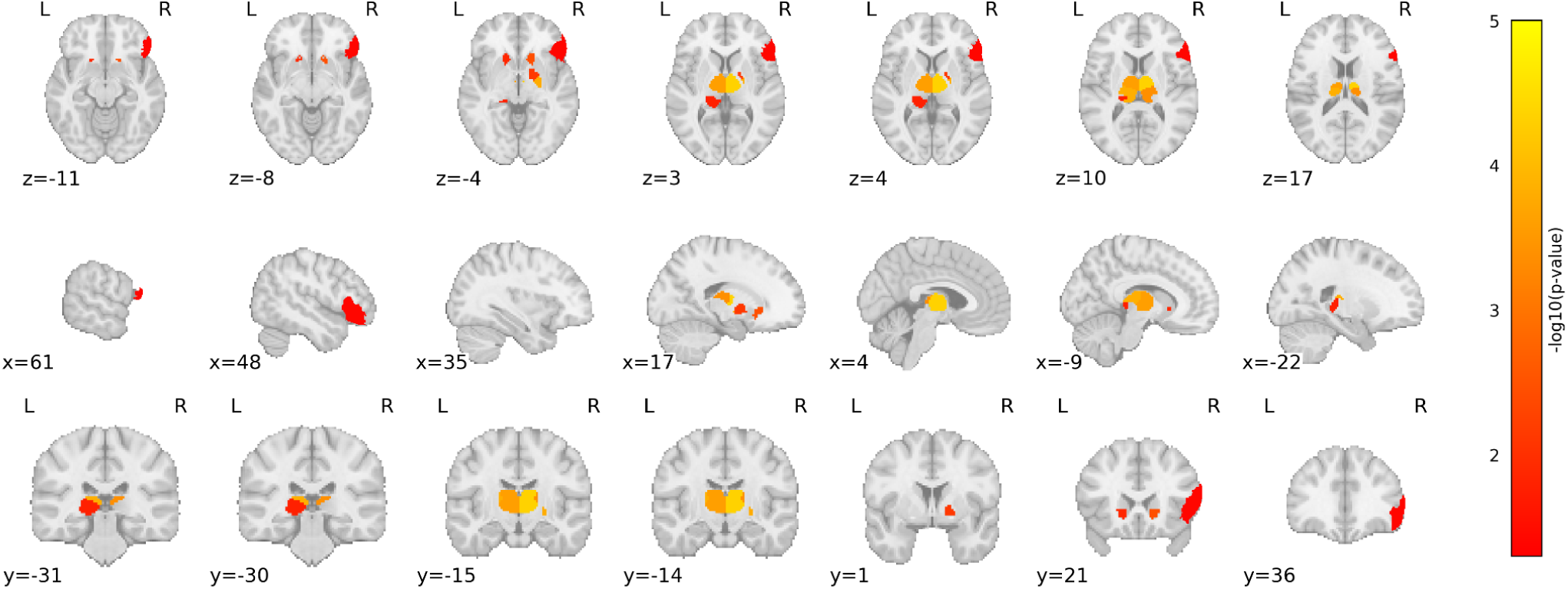
Brain regions with significantly different connectivity patterns between RS from healthy controls and RS from patients with early psychosis, as identified by OSCAR (L = left, R = right, x = sagittal, y = coronal, z = axial).

## 4. Discussion

We here proposed and validated a new method OSCAR to identify multivariate group differences in BOLD connectivity patterns of brain regions. We first showed that OSCAR provides novel and interpretable insights by applying it to FC data from three different tasks across three healthy cohorts. Moreover, OSCAR detected connectivity alterations in regions previously established as task-relevant in the literature, which perMANOVA failed to identify. These results were further supported by the observation that OSCAR-identified regions were often spatially closer to established task-specific activation findings than those identified by perMANOVA. Additionally, in a sample with early psychosis patients, OSCAR likewise identified connectivity differences in disease-related regions that are known to be affected in early psychosis not identified by a perMANOVA.

### 4.1 GStroop Task

Both OSCAR and perMANOVA identified several regions that are known to be involved in the processing of the (gender-) Stroop-task such as the precuneus (Schaefer: RH_ContC_pCun_2, HO: Precuneous Cortex) (Harrison et al., 2005; Milham et al., 2003), the posterior cingulate gyrus (Schaefer: RH_DefaultA_pCunPCC_1, HO: Cingulate Gyrus, posterior division) (Harrison et al., 2005; Milham et al., 2003), the precentral gyrus (Schaefer: RH_SalVentAttnA_PrC_1, HO: Precentral Gyrus) (Tang et al., 2006) and the lateral prefrontal cortex (Schaefer: LH_SalVentAttnB_PFCl_1, HO: Frontal Pole) (Wang et al., 2023). In addition, OSCAR detected connectivity differences in 78 regions that were not detected by perMANOVA (Fig. 2). Notably, only OSCAR identified the insula (Schaefer: RH_SalVentAttnA_Ins_2 and RH_SalVentAttnA_FrOper_1, HO: Insular Cortex) and the anterior cingulate cortex (Schaefer: RH_ContA_Cingm_1 and LH_ContA_Cingm_1, HO: Cingulate Gyrus, anterior division), both involved in conflict monitoring during the stroop task (Dragan et al., 2022; Huang et al., 2020; Swick and Jovanovic, 2002). OSCAR also detected several other regions known to be involved in the stroop task such as the caudate (Tian: aCAU-lh, HO: Left Caudate) (Ali et al., 2010), parts of the thalamus (Tian: THA-VA-lh, THA-DP-rh and THA-VP-lh, HO: Left Thalamus, Right Thalamus) (Parris et al., 2019) and inferior frontal gyrus (Schaefer: RH_ContA_PFCl_1 and LH_ContA_PFClv_1 HO: Inferior Frontal Gyrus, pars opercularis and Inferior Frontal Gyrus, pars triangularis) (Huang et al., 2020). Spatial correspondence with the peak activations in the whole-brain group-level results from the AOMIC dataset analysis (Snoek et al., 2021) showed that the regions identified by OSCAR were significantly closer to the positive contrast (incorrect > correct) than those detected by the perMANOVA (Fig. 2d).

### 4.2 OLM Task

Applied to data from the OLM-task, both methods identified connectivity differences in several regions (OSCAR 48, perMANOVA 37). Among these were several regions that are a part of the ventral visual stream that is necessary for object identification (Mishkin et al., 1983; Rolls et al., 2024) and play a prominent role during the OLM-task. Further, OSCAR and perMANOVA identified the precuneus (Schaefer: LH_DefaultA_pCunPCC_1 and RH_ContC_pCun_2, HO: Precuneous Cortex), the retrospenial cortex (Schaefer: RH_DefaultC_Rsp_1, HO: Precuneous Cortex), the middle frontal gyrus (Schaefer: LH_ContB_PFCl_1, RH_DefaultA_PFCd_1, HO: Middle Frontal Gyrus), the insula cortex (Schaefer: RH_SalVentAttnB_Ins_2, HO: Frontal Opercular Cortex/Insular Cortex) and the precentral gyrus (Schaefer: LH_SomMotB_Cent_2, HO: Precentral Gyrus) all of which are known to be related to visual spatial learning (Bogler et al., 2024; Gillis et al., 2016a; Postma et al., 2008). In addition, both methods identified the lateral occipital complex, (Schaefer: RH_VisPeri_ExStrSup_3, RH_VisCent_ExStr_5 LH_VisCent_ExStr_5 LH_DefaultC_IPL_1, LH_DefaultA_IPL_1 HO: Lateral Occipital Cortex inferior and superior division) which is tied to object recognition, object form perception and object naming (Gillis et al., 2016a; Petersson et al., 2001).

Only OSCAR identified connectivity changes in further regions that have been indicated to be involved in object location and memory tasks by Gillis et al such as the bilateral Pallidum (Tian: pGP-lh, pGP-rh, HO: left & right Pallidum), and the left thalamus (Tian: THA-VP-lh, HO: Left Thalamus) (Gillis et al., 2016b)). In addition OSCAR identified the frontal orbital cortex (Schaefer: LH_DefaultB_PFCv_3, HO: Frontal Orbital Cortex), which might reflect feedback-based learning and decision-making (Wallis, 2007) and parts of the left hemisphere inferior and middle temporo-occipital cortices (e.g. Schaefer: LH_ContA_Temp_1 / LH_ContB_Temp_1, HO: Middle Temporal Gyrus, temporo-occipital part / Inferior Temporal Gyrus, temporo-occipital part) which are involved in object processing (Gillis et al., 2016b; Rolls et al., 2024) (Fig. 3). Notably, the regions identified by OSCAR largely overlap with a general linear model analysis from a recent publication on the same data by Abdelmotaleb et al (Abdelmotaleb et al., 2025). Compared with the significant clusters of task activity found by Abdelmotaleb et al., OSCAR-identified regions were closer on average but not statistically significantly closer than perMANOVA (Fig. 4c and 4d).

### 4.3 NWL Task

For the Novel Word Learning task, both methods detected the left superior temporal gyrus (Schaefer: LH_DefaultB_Temp_3, HO: superior temporal gyrus, anterior division), which is a core region for phonological processing, speech perception, and lexical access (Mesgarani et al., 2014; Price, 2000). In addition, both methods identified changes in the extra-striate superior (Schaefer: RH_VisPeri_ExStrSup_3, HO: Lateral Occipital Cortex, superior division) which might reflect the visual component of the task stimulus.

OSCAR additionally identified connectivity changes in the medial prefrontal cortex (Schaefer: LH_DefaultA_PFCm_2, HO: Frontal Pole/Frontal Medial Cortex), which supports the integration of new information into existing schemas, as required when linking pseudowords to familiar objects in the NWL task (Preston and Eichenbaum, 2013; Schlichting and Preston, 2015; Van Kesteren et al., 2013) and bilateral fusiform gyrus (Schaefer: LH_VisCent_ExStr_1 and RH_VisCent_ExStr_1, HO: Occipital Fusiform Gyrus), which has been demonstrated to be involved in lexical learning (Tagarelli et al., 2019). Further, OSCAR detected changes in the globus pallidus (Tian: pGP-lh and pGP-rh, HO: Left Pallidum and Right Pallidum) which is tied to language functions (Tomasi and Volkow, 2012), the orbito-frontal cortex (Schaefer: RH_LimbicB_OFC_4, HO: Frontal Pole) which showed a significant main effect of learning stage during the NWL-task in (Sliwinska et al., 2017) (Fig.5).

OSCAR also detected changes in various visual regions, where connectivity changes are likely induced by the colored pictures of objects that represent pseudowords during the learning stage of the task. The regions that were identified by OSCAR partly match findings from an activation study on the same data by Kocataş et al (Kocataş et al., 2025). Both studies detected differences during the learning task in bilateral occipito-temporal regions and temporal regions. However, only OSCAR identified regions in the frontal pole.

Notably, OSCAR did not detect changes in the inferior frontal gyrus (IFG), left supplementary area or Insula, all of which have reliably shown strong effects in previous activation studies (Kocataş et al., 2025; Sliwinska et al., 2017; Tagarelli et al., 2019). To further investigate this discrepancy, we conducted an additional parcel-wise GLM analysis on the same dataset (Supplementary Table 9). This analysis revealed significant task-evoked activity in 20 regions, among them the left IFG, left supplementary motor area, and insula. Possibly, the task-evoked activity in these regions does not translate into widespread connectivity reconfiguration. Instead, changes may be confined to a few specific connections, and thus remain undetected when focusing on differences in region-to-whole-brain connectivity profiles as is done by OSCAR. Further, conventional task-based fMRI analyses model the time-locked structure of stimulus presentation using event-related or block-design regressors, enabling detection of transient, spatially localized responses. Such effects may be present only briefly and, when connectivity is aggregated across time, fail to produce sustained or distributed changes in region-to-whole-brain connectivity detectable by OSCAR.

Only perMANOVA detected the inferior parietal lobule (Schaefer: RH_SalVentAttnB_IPL_1, HO: Supramarginal Gyrus, posterior division) and the insula (Schaefer: RH_SalVentAttnA_Ins_2, HO: Planum Polare/Insular Cortex), both of which are tied to the NWL-task (Sliwinska et al., 2017). Compared to results from a meta-analysis on lexical learning, both methods achieved similar spatial correspondence to the peaks identified by an activation likelihood estimation meta-analysis (Tagarelli et al., 2019) (Fig. 6c).

### 4.4 Early psychosis

Applied to data from patients with early psychosis, both methods identified three thalamic regions, the dorsal anterior subregion on both hemispheres and the ventral anterior subregion on the left hemisphere. OSCAR additionally identified differences in more parts of the thalamus, the nucleus accumbens, the globus pallidus as well as the ventromedial prefrontal cortex, all of which are known to be affected in psychosis (Bois et al., 2015; Jensen et al., 2024; Mamah et al., 2007; Qi et al., 2023) (Fig. 7). These results align with the previous literature as the thalamus has long been associated with psychosis and is believed to play a key role in its pathophysiology, affecting sensory, cognitive, and sleep processes (Alemán-Gómez et al., 2023; Onofrj et al., 2023).

### 4.5 Limitations

Several limitations should be considered when interpreting our findings. First, several regions previously implicated in specific tasks based on activation analysis (e.g. the IFG for the NWL task) were not detected either by OSCAR or perMANOVA, suggesting task-related activation of these regions may not coincide with large-scale connectivity reorganization. Instead, changes may be confined to a few specific connections, and thus remain undetected when focusing on differences in region-to-whole-brain connectivity profiles as done in our analysis. However, we do not see OSCAR as an alternative to traditional activation analyses, but rather as providing complementary insights. Furthermore, OSCAR currently uses whole brain connectivity profiles which may restrict its sensitivity when the majority of connectivity across states remain unchanged. Restricting OSCAR to refined connectivity profiles could increase its sensitivity. Second, the datasets analyzed in this study were acquired across multiple sites, introducing potential variability related to scanner hardware, acquisition protocols and other factors (Yan et al., 2013). Such heterogeneity may contribute to systematic differences or subtle site effects. While the use of a standardized preprocessing pipeline (fMRIPrep) helps to reduce such variability, residual site effects cannot be fully ruled out. However, the results here were largely consistent with prior findings reported in the literature. Third, this study employed the Schaefer 200 regions × 17 networks cortical parcellation complemented with subcortical regions from the Melbourne atlas. The choice of parcellation can influence the sensitivity and specificity of detecting FC differences, as the spatial resolution and network definitions determine how finely regional signals are represented. Larger parcels may pool signals across functionally heterogeneous subregions, potentially masking localized effects, whereas smaller parcels increase sensitivity to subtle differences but are more vulnerable to noise. As a result, the specific parcellation scheme employed may bias which regional effects are detectable, and future work should analyze the influence of using different granularities of a given atlas as well as restricting connectivity profiles to networks instead of using the whole brain. Finally, the reliability of FC is strongly dependent on the length of a sequence. The connectomes used to train OSCAR were derived from scans with a length of 6-8 minutes. Although this is a commonly used scan length, previous research has indicated that reliability can be greatly improved by using sequences of 9 to 13 minutes or longer (Birn et al., 2013). Thus, the performance of OSCAR may further benefit from training on data acquired with longer scan durations.

## 5. Summary and conclusion

Taken together, we demonstrated here that OSCAR can identify regions where FC profile deviates due to the task engagement or a disease state. Importantly, OSCAR extended the findings of a standard perMANOVA by revealing effects in additional condition-related regions, suggesting that OSCAR is more sensitive to the multivariate effects induced by different conditions. OSCAR captures the multivariate connectivity pattern of a given region during the reference state and then identifies deviations due to a condition. In contrast, perMANOVA tests whether the mean connectivity pattern (the “centroid”) between the two groups differs spatially (Anderson, 2001). Thus perMANOVA is sensitive to the dispersion of points but it can not detect deviations in the distribution of the data (Anderson, 2001).

Therefore, OSCAR might be more sensitive to subtle differences in FC patterns induced by a task or disease. Beyond its empirical performance, OSCAR also provides a conceptual contribution to the analysis of condition-related FC. Traditional task-based activation analyses identify localized changes in BOLD amplitude, and thus reveal where neural responses increase or decrease during a task (Friston et al., 1994). However, activation maps do not capture whether the pattern of connectivity between a region and the rest of the brain changes (Friston, 2011). OSCAR quantifies how a region’s multivariate connectivity profile deviates across states, allowing researchers to detect condition-relevant connectivity reconfigurations which might not be accompanied by local activation. Conceptually, OSCAR enables a middle ground between univariate activation mapping and unconstrained multivariate decomposition. OSCAR can be used to (1) identify relevant regions whose connectivity reorganizes due to cognitive processing, (2) detect clinically relevant deviations in patient groups, (3) generate hypotheses about compensatory or maladaptive network reconfiguration, and (4) provide region-level readouts that are directly comparable across tasks, datasets, and cohorts. This region-centered framing of OSCAR can add value to analysis of data from causal perturbation approaches like invasive or non-invasive brain stimulation approaches (e.g., transcranial magnetic stimulation (Siebner et al., 2022), deep brain stimulation (Krauss et al., 2021) or transcranial direct current stimulation (Meinzer et al., 2024), where localized interventions shift the embedding of both targeted and downstream regions and thereby provide a principled way to map network effects.

Together, these findings demonstrate that OSCAR is able to identify multivariate group differences in BOLD connectivity patterns of brain regions. Compared to perMANOVA, OSCAR showed an advantage in identifying condition-related regions across multiple experimental tasks as well as in a patient cohort. We anticipate that OSCAR will serve as a valuable tool for researchers aiming to map condition-specific changes in FC, whether in task paradigms, in clinical populations, or in perturbation approaches like invasive or non-invasive brain stimulation.

## Funding

This work was supported by the eBRAIN-Health project. eBRAIN-Health has received funding from the European Union’s HorizonEurope research and innovation programme under grant agreement No 101058516 and HORIZON.1.3, grant no. 101147319. This work was also funded by the German Research Foundation (Research Unit5429/1 [467143400]; CRC1315-B03 [327654276]; FL379/22-1; FL379/26-1; FL379/34-1; FL379/35-1; FL379/37-1; ME3161/5-1; ME3161/6-1).

## Declaration of interests

The authors declare that they have no known competing financial interests or personal relationships that could have appeared to influence the work reported in this paper.

## Declaration of generative AI and AI-assisted technologies in the manuscript preparation process

During the preparation of this work the author(s) used ChatGPT in order to draft text / coding. After using this tool/service, the author(s) reviewed and edited the content as needed and take(s) full responsibility for the content of the published article.

## Notes

### Competing Interest Statement

The authors have declared no competing interest.

### Summary of Updates

Changed author name also outside of manuscript

## References

1. Abdelmotaleb, M., Niemann, F., Kocataş, H., Caisachana Guevara, L.M., Shahbabaie, A., Malinowski, R., Riemann, S., Fromm, A.E., Hayek, D., Antonenko, D., Meinzer, M., Flöel, A., 2025. Identification of Reliable Target Brain Regions for Enhancing Object–Location Memory by Brain Stimulation. Brain and Behavior 15, e70658. 10.1002/brb3.70658

2. Alemán-Gómez, Y., Baumgartner, T., Klauser, P., Cleusix, M., Jenni, R., Hagmann, P., Conus, P., Do, K.Q., Bach Cuadra, M., Baumann, P.S., Steullet, P., 2023. Multimodal Magnetic Resonance Imaging Depicts Widespread and Subregion Specific Anomalies in the Thalamus of Early-Psychosis and Chronic Schizophrenia Patients. Schizophrenia Bulletin 49, 196–207. 10.1093/schbul/sbac113

3. Ali, N., Green, D.W., Kherif, F., Devlin, J.T., Price, C.J., 2010. The role of the left head of caudate in suppressing irrelevant words. J Cogn Neurosci 22, 2369–2386. 10.1162/jocn.2009.21352

4. Anderson, M.J., 2001. A new method for non-parametric multivariate analysis of variance. Austral Ecology 26, 32–46. 10.1111/j.1442-9993.2001.01070.pp.x

5. Avants, B.B., Tustison, N.J., Song, G., Cook, P.A., Klein, A., Gee, J.C., 2011. A reproducible evaluation of ANTs similarity metric performance in brain image registration. NeuroImage 54, 2033–2044. 10.1016/j.neuroimage.2010.09.025

6. Birn, R.M., Molloy, E.K., Patriat, R., Parker, T., Meier, T.B., Kirk, G.R., Nair, V.A., Meyerand, M.E., Prabhakaran, V., 2013. The effect of scan length on the reliability of resting-state fMRI connectivity estimates. NeuroImage 83, 550–558. 10.1016/j.neuroimage.2013.05.099

7. Bogler, C., Zangrossi, A., Miller, C., Sartori, G., Haynes, J., 2024. Have you been there before? Decoding recognition of spatial scenes from FMRI signals in precuneus. Human Brain Mapping 45, e26690. 10.1002/hbm.26690

8. Bois, C., Levita, L., Ripp, I., Owens, D.C.G., Johnstone, E.C., Whalley, H.C., Lawrie, S.M., 2015. Hippocampal, amygdala and nucleus accumbens volume in first-episode schizophrenia patients and individuals at high familial risk: A cross-sectional comparison. Schizophrenia Research 165, 45–51. 10.1016/j.schres.2015.03.024

9. Bressler, S.L., Menon, V., 2010. Large-scale brain networks in cognition: emerging methods and principles. Trends in Cognitive Sciences 14, 277–290. 10.1016/j.tics.2010.04.004

10. Cattarinussi, G., Grimaldi, D.A., Aarabi, M.H., Sambataro, F., 2024. Static and Dynamic Dysconnectivity in Early Psychosis: Relationship With Symptom Dimensions. Schizophrenia Bulletin 51, 120–132. 10.1093/schbul/sbae142

11. Desikan, R.S., Ségonne, F., Fischl, B., Quinn, B.T., Dickerson, B.C., Blacker, D., Buckner, R.L., Dale, A.M., Maguire, R.P., Hyman, B.T., Albert, M.S., Killiany, R.J., 2006. An automated labeling system for subdividing the human cerebral cortex on MRI scans into gyral based regions of interest. NeuroImage 31, 968–980. 10.1016/j.neuroimage.2006.01.021

12. D’Esposito, M., Postle, B.R., Rypma, B., 2000. Prefrontal cortical contributions to working memory: evidence from event-related fMRI studies. Exp Brain Res 133, 3–11. 10.1007/s002210000395

13. Dong, D., Wang, Y., Chang, X., Luo, C., Yao, D., 2018. Dysfunction of Large-Scale Brain Networks in Schizophrenia: A Meta-analysis of Resting-State Functional Connectivity. Schizophrenia Bulletin 44, 168–181. 10.1093/schbul/sbx034

14. Dragan, W.Ł., Sokołowski, A., Folkierska-Żukowska, M., 2022. Temperament and neural activation during the affective Stroop task: A functional connectivity study. Personality and Individual Differences 186, 111385. 10.1016/j.paid.2021.111385

15. Dunn, O.J., 1961. Multiple Comparisons among Means. Journal of the American Statistical Association 56, 52–64. 10.1080/01621459.1961.10482090

16. Esteban, O., Markiewicz, C.J., Blair, R.W., Moodie, C.A., Isik, A.I., Erramuzpe, A., Kent, J.D., Goncalves, M., DuPre, E., Snyder, M., Oya, H., Ghosh, S.S., Wright, J., Durnez, J., Poldrack, R.A., Gorgolewski, K.J., 2019. fMRIPrep: a robust preprocessing pipeline for functional MRI. Nat Methods 16, 111–116. 10.1038/s41592-018-0235-4

17. Etzel, J.A., Zacks, J.M., Braver, T.S., 2013. Searchlight analysis: Promise, pitfalls, and potential. NeuroImage 78, 261–269. 10.1016/j.neuroimage.2013.03.041

18. Fisher, R.A., 1915. FREQUENCY DISTRIBUTION OF THE VALUES OF THE CORRELATION COEFFIENTS IN SAMPLES FROM AN INDEFINITELY LARGE POPU;ATION. Biometrika 10, 507–521. 10.1093/biomet/10.4.507

19. Friston, K., 2002. Functional integration and inference in the brain. Progress in Neurobiology 68, 113–143. 10.1016/S0301-0082(02)00076-X

20. Friston, K.J., 2011. Functional and Effective Connectivity: A Review. Brain Connectivity 1, 13–36. 10.1089/brain.2011.0008

21. Friston, K.J., Holmes, A.P., Worsley, K.J., Poline, J.-P., Frith, C.D., Frackowiak, R.S.J., 1994. Statistical parametric maps in functional imaging: A general linear approach. Human Brain Mapping 2, 189–210. 10.1002/hbm.460020402

22. Gilead, M., Liberman, N., Maril, A., 2014. From mind to matter: neural correlates of abstract and concrete mindsets. Social Cognitive and Affective Neuroscience 9, 638–645. 10.1093/scan/nst031

23. Gillis, M.M., Garcia, S., Hampstead, B.M., 2016a. Working memory contributes to the encoding of object location associations: Support for a 3-part model of object location memory. Behavioural Brain Research 311, 192–200. 10.1016/j.bbr.2016.05.037

24. Gillis, M.M., Garcia, S., Hampstead, B.M., 2016b. Working memory contributes to the encoding of object location associations: Support for a 3-part model of object location memory. Behavioural Brain Research 311, 192–200. 10.1016/j.bbr.2016.05.037

25. Glasser, M.F., Sotiropoulos, S.N., Wilson, J.A., Coalson, T.S., Fischl, B., Andersson, J.L., Xu, J., Jbabdi, S., Webster, M., Polimeni, J.R., Van Essen, D.C., Jenkinson, M., 2013. The minimal preprocessing pipelines for the Human Connectome Project. NeuroImage 80, 105–124. 10.1016/j.neuroimage.2013.04.127

26. Greve, D.N., Fischl, B., 2009. Accurate and robust brain image alignment using boundary-based registration. NeuroImage 48, 63–72. 10.1016/j.neuroimage.2009.06.060

27. Harrison, B.J., Shaw, M., Yücel, M., Purcell, R., Brewer, W.J., Strother, S.C., Egan, G.F., Olver, J.S., Nathan, P.J., Pantelis, C., 2005. Functional connectivity during Stroop task performance. NeuroImage 24, 181–191. 10.1016/j.neuroimage.2004.08.033

28. Huang, S., De Brigard, F., Cabeza, R., Davis, S.W., 2024. Connectivity analyses for task-based fMRI. Physics of Life Reviews 49, 139–156. 10.1016/j.plrev.2024.04.012

29. Huang, Y., Su, L., Ma, Q., 2020. The Stroop effect: An activation likelihood estimation meta-analysis in healthy young adults. Neuroscience Letters 716, 134683. 10.1016/j.neulet.2019.134683

30. Jacobs, G.R., Coleman, M.J., Lewandowski, K.E., Pasternak, O., Cetin-Karayumak, S., Mesholam-Gately, R.I., Wojcik, J., Kennedy, L., Knyazhanskaya, E., Reid, B., Swago, S., Lyons, M.G., Rizzoni, E., John, O., Carrington, H., Kim, N., Kotler, E., Veale, S., Haidar, A., Prunier, N., Haaf, M., Levitt, J.J., Seitz-Holland, J., Rathi, Y., Kubicki, M., Keshavan, M.S., Holt, D.J., Seidman, L.J., Öngür, D., Breier, A., Bouix, S., Shenton, M.E., 2024. An Introduction to the Human Connectome Project for Early Psychosis. Schizophrenia Bulletin sbae123. 10.1093/schbul/sbae123

31. Jenkinson, M., Bannister, P., Brady, M., Smith, S., 2002. Improved Optimization for the Robust and Accurate Linear Registration and Motion Correction of Brain Images. NeuroImage 17, 825–841. 10.1006/nimg.2002.1132

32. Jensen, K.M., Calhoun, V.D., Fu, Z., Yang, K., Faria, A.V., Ishizuka, K., Sawa, A., Andrés-Camazón, P., Coffman, B.A., Seebold, D., Turner, J.A., Salisbury, D.F., Iraji, A., 2024. A whole-brain neuromark resting-state fMRI analysis of first-episode and early psychosis: Evidence of aberrant cortical-subcortical-cerebellar functional circuitry. NeuroImage: Clinical 41, 103584. 10.1016/j.nicl.2024.103584

33. Kanwisher, N., 2010. Functional specificity in the human brain: A window into the functional architecture of the mind. Proc. Natl. Acad. Sci. U.S.A. 107, 11163–11170. 10.1073/pnas.1005062107

34. Kocataş, H., Abdelmotaleb, M., Caisachana Guevara, L.M., Niemann, F., Shahbabaie, A., Malinowski, R., Riemann, S., Hayek, D., Antonenko, D., Rodriguez-Fornells, A., Flöel, A., Meinzer, M., 2025. Functionally Relevant and Reliable Brain Stimulation Targets for Enhancement of Novel Word-Learning. 10.1101/2025.11.04.686295

35. Kolskår, K.K., Alnæs, D., Kaufmann, T., Richard, G., Sanders, A.-M., Ulrichsen, K.M., Moberget, T., Andreassen, O.A., Nordvik, J.E., Westlye, L.T., 2018. Key Brain Network Nodes Show Differential Cognitive Relevance and Developmental Trajectories during Childhood and Adolescence. eNeuro 5, ENEURO.0092-18.2018. 10.1523/ENEURO.0092-18.2018

36. Krauss, J.K., Lipsman, N., Aziz, T., Boutet, A., Brown, P., Chang, J.W., Davidson, B., Grill, W.M., Hariz, M.I., Horn, A., Schulder, M., Mammis, A., Tass, P.A., Volkmann, J., Lozano, A.M., 2021. Technology of deep brain stimulation: current status and future directions. Nat Rev Neurol 17, 75–87. 10.1038/s41582-020-00426-z

37. Kriegeskorte, N., 2008. Representational similarity analysis – connecting the branches of systems neuroscience. Front. Sys. Neurosci. 10.3389/neuro.06.004.2008

38. Makary, M.M., Eun, S., Soliman, R.S., Mohamed, A.Z., Lee, J., Park, K., 2017. Functional topography of the primary motor cortex during motor execution and motor imagery as revealed by functional MRI. NeuroReport 28, 731–738. 10.1097/WNR.0000000000000825

39. Mamah, D., Wang, L., Barch, D., De Erausquin, G.A., Gado, M., Csernansky, J.G., 2007. Structural analysis of the basal ganglia in schizophrenia. Schizophrenia Research 89, 59–71. 10.1016/j.schres.2006.08.031

40. Marquand, A.F., Rezek, I., Buitelaar, J., Beckmann, C.F., 2016. Understanding Heterogeneity in Clinical Cohorts Using Normative Models: Beyond Case-Control Studies. Biological Psychiatry 80, 552–561. 10.1016/j.biopsych.2015.12.023

41. Mckeown, M.J., Makeig, S., Brown, G.G., Jung, T.-P., Kindermann, S.S., Bell, A.J., Sejnowski, T.J., 1998. Analysis of fMRI data by blind separation into independent spatial components. Hum. Brain Mapp. 6, 160–188. 10.1002/(SICI)1097-0193(1998)6:3%253C160::AID-HBM5%253E3.0.CO;2-1

42. Meinzer, M., Shahbabaie, A., Antonenko, D., Blankenburg, F., Fischer, R., Hartwigsen, G., Nitsche, M.A., Li, S.-C., Thielscher, A., Timmann, D., Waltemath, D., Abdelmotaleb, M., Kocataş, H., Caisachana Guevara, L.M., Batsikadze, G., Grundei, M., Cunha, T., Hayek, D., Turker, S., Schlitt, F., Shi, Y., Khan, A., Burke, M., Riemann, S., Niemann, F., Flöel, A., 2024. Investigating the neural mechanisms of transcranial direct current stimulation effects on human cognition: current issues and potential solutions. Front. Neurosci. 18, 1389651. 10.3389/fnins.2024.1389651

43. Mesgarani, N., Cheung, C., Johnson, K., Chang, E.F., 2014. Phonetic Feature Encoding in Human Superior Temporal Gyrus. Science 343, 1006–1010. 10.1126/science.1245994

44. Milham, M.P., Banich, M.T., Barad, V., 2003. Competition for priority in processing increases prefrontal cortex’s involvement in top-down control: an event-related fMRI study of the stroop task. Cognitive Brain Research 17, 212–222. 10.1016/S0926-6410(03)00108-3

45. Mishkin, M., Ungerleider, L.G., Macko, K.A., 1983. Object vision and spatial vision: two cortical pathways. Trends in Neurosciences 6, 414–417. 10.1016/0166-2236(83)90190-X

46. Murphy, A.C., Gu, S., Khambhati, A.N., Wymbs, N.F., Grafton, S.T., Satterthwaite, T.D., Bassett, D.S., 2016. Explicitly Linking Regional Activation and Function Connectivity: Community Structure of Weighted Networks with Continuous Annotation. 10.48550/ARXIV.1611.07962

47. Nieto-Castanon, A., 2022. Brain-wide connectome inferences using functional connectivity MultiVariate Pattern Analyses (fc-MVPA). PLoS Comput Biol 18, e1010634. 10.1371/journal.pcbi.1010634

48. Onofrj, M., Russo, M., Delli Pizzi, S., De Gregorio, D., Inserra, A., Gobbi, G., Sensi, S.L., 2023. The central role of the Thalamus in psychosis, lessons from neurodegenerative diseases and psychedelics. Transl Psychiatry 13, 384. 10.1038/s41398-023-02691-0

49. Parris, B.A., Wadsley, M.G., Hasshim, N., Benattayallah, A., Augustinova, M., Ferrand, L., 2019. An fMRI Study of Response and Semantic Conflict in the Stroop Task. Front. Psychol. 10, 2426. 10.3389/fpsyg.2019.02426

50. Petersson, K.M., Sandblom, J., Gisselgård, J., Ingvar, M., 2001. Learning related modulation of functional retrieval networks in man. Scandinavian J Psychology 42, 197–216. 10.1111/1467-9450.00231

51. Postma, A., Kessels, R., Vanasselen, M., 2008. How the brain remembers and forgets where things are: The neurocognition of object–location memory. Neuroscience & Biobehavioral Reviews 32, 1339–1345. 10.1016/j.neubiorev.2008.05.001

52. Preston, A.R., Eichenbaum, H., 2013. Interplay of Hippocampus and Prefrontal Cortex in Memory. Current Biology 23, R764–R773. 10.1016/j.cub.2013.05.041

53. Price, C.J., 2000. The anatomy of language: contributions from functional neuroimaging. Journal of Anatomy 197, 335–359. 10.1046/j.1469-7580.2000.19730335.x

54. Qi, W., Wen, Z., Chen, J., Capichioni, G., Ando, F., Chen, Z.S., Wang, J., Yoncheva, Y., Castellanos, F.X., Milad, M., Goff, D.C., 2023. Aberrant resting-state functional connectivity of the globus pallidus interna in first-episode schizophrenia. Schizophrenia Research 261, 100–106. 10.1016/j.schres.2023.09.018

55. Rolls, E.T., Zhang, R., Deco, G., Vatansever, D., Feng, J., 2024. Selective Brain Activations and Connectivities Related to the Storage and Recall of Human Object-Location, Reward-Location, and Word-Pair Episodic Memories. Human Brain Mapping 45, e70056. 10.1002/hbm.70056

56. Rutherford, S., Barkema, P., Tso, I.F., Sripada, C., Beckmann, C.F., Ruhe, H.G., Marquand, A.F., 2023. Evidence for embracing normative modeling. eLife 12, e85082. 10.7554/eLife.85082

57. Salimi-Khorshidi, G., Douaud, G., Beckmann, C.F., Glasser, M.F., Griffanti, L., Smith, S.M., 2014. Automatic denoising of functional MRI data: Combining independent component analysis and hierarchical fusion of classifiers. NeuroImage 90, 449–468. 10.1016/j.neuroimage.2013.11.046

58. Satake, T., Taki, A., Kasahara, K., Yoshimaru, D., Tsurugizawa, T., 2024. Comparison of local activation, functional connectivity, and structural connectivity in the N-back task. Front Neurosci 18, 1337976. 10.3389/fnins.2024.1337976

59. Schaefer, A., Kong, R., Gordon, E.M., Laumann, T.O., Zuo, X.-N., Holmes, A.J., Eickhoff, S.B., Yeo, B.T.T., 2018. Local-Global Parcellation of the Human Cerebral Cortex from Intrinsic Functional Connectivity MRI. Cerebral Cortex 28, 3095–3114. 10.1093/cercor/bhx179

60. Schlichting, M.L., Preston, A.R., 2015. Memory integration: neural mechanisms and implications for behavior. Current Opinion in Behavioral Sciences 1, 1–8. 10.1016/j.cobeha.2014.07.005

61. Schölkopf, B., Williamson, R.C., Smola, A., Shawe-Taylor, J., Platt, J., 1999. Support Vector Method for Novelty Detection, in: Solla, S., Leen, T., Müller, K. (Eds.), Advances in Neural Information Processing Systems. MIT Press.

62. Segal, A., Parkes, L., Aquino, K., Kia, S.M., Wolfers, T., Franke, B., Hoogman, M., Beckmann, C.F., Westlye, L.T., Andreassen, O.A., Zalesky, A., Harrison, B.J., Davey, C.G., Soriano-Mas, C., Cardoner, N., Tiego, J., Yücel, M., Braganza, L., Suo, C., Berk, M., Cotton, S., Bellgrove, M.A., Marquand, A.F., Fornito, A., 2023. Regional, circuit and network heterogeneity of brain abnormalities in psychiatric disorders. Nat Neurosci 26, 1613–1629. 10.1038/s41593-023-01404-6

63. Shine, J.M., Bissett, P.G., Bell, P.T., Koyejo, O., Balsters, J.H., Gorgolewski, K.J., Moodie, C.A., Poldrack, R.A., 2016. The Dynamics of Functional Brain Networks: Integrated Network States during Cognitive Task Performance. Neuron 92, 544–554. 10.1016/j.neuron.2016.09.018

64. Siebner, H.R., Funke, K., Aberra, A.S., Antal, A., Bestmann, S., Chen, R., Classen, J., Davare, M., Di Lazzaro, V., Fox, P.T., Hallett, M., Karabanov, A.N., Kesselheim, J., Beck, M.M., Koch, G., Liebetanz, D., Meunier, S., Miniussi, C., Paulus, W., Peterchev, A.V., Popa, T., Ridding, M.C., Thielscher, A., Ziemann, U., Rothwell, J.C., Ugawa, Y., 2022. Transcranial magnetic stimulation of the brain: What is stimulated? – A consensus and critical position paper. Clinical Neurophysiology 140, 59–97. 10.1016/j.clinph.2022.04.022

65. Sliwinska, M.W., Violante, I.R., Wise, R.J.S., Leech, R., Devlin, J.T., Geranmayeh, F., Hampshire, A., 2017. Stimulating Multiple-Demand Cortex Enhances Vocabulary Learning. J. Neurosci. 37, 7606–7618. 10.1523/JNEUROSCI.3857-16.2017

66. Snoek, L., van der Miesen, M.M., Beemsterboer, T., van der Leij, A., Eigenhuis, A., Steven Scholte, H., 2021. The Amsterdam Open MRI Collection, a set of multimodal MRI datasets for individual difference analyses. Sci Data 8, 85. 10.1038/s41597-021-00870-6

67. Swick, D., Jovanovic, J., 2002. Anterior cingulate cortex and the Stroop task: neuropsychological evidence for topographic specificity. Neuropsychologia 40, 1240–1253. 10.1016/S0028-3932(01)00226-3

68. Tagarelli, K.M., Shattuck, K.F., Turkeltaub, P.E., Ullman, M.T., 2019. Language learning in the adult brain: A neuroanatomical meta-analysis of lexical and grammatical learning. NeuroImage 193, 178–200. 10.1016/j.neuroimage.2019.02.061

69. Tang, J., Critchley, H.D., Glaser, D.E., Dolan, R.J., Butterworth, B., 2006. Imaging informational conflict: a functional magnetic resonance imaging study of numerical stroop. J Cogn Neurosci 18, 2049–2062. 10.1162/jocn.2006.18.12.2049

70. Tian, Y., Margulies, D.S., Breakspear, M., Zalesky, A., 2020. Topographic organization of the human subcortex unveiled with functional connectivity gradients. Nat Neurosci 23, 1421–1432. 10.1038/s41593-020-00711-6

71. Tomasi, D., Volkow, N.D., 2012. Resting functional connectivity of language networks: characterization and reproducibility. Mol Psychiatry 17, 841–854. 10.1038/mp.2011.177

72. Van Kesteren, M.T.R., Beul, S.F., Takashima, A., Henson, R.N., Ruiter, D.J., Fernández, G., 2013. Differential roles for medial prefrontal and medial temporal cortices in schema-dependent encoding: From congruent to incongruent. Neuropsychologia 51, 2352–2359. 10.1016/j.neuropsychologia.2013.05.027

73. Wallis, J.D., 2007. Orbitofrontal Cortex and Its Contribution to Decision-Making. Annu. Rev. Neurosci. 30, 31–56. 10.1146/annurev.neuro.30.051606.094334

74. Wang, J.-X., Li, Y., Mu, Y., Zhuang, J.-Y., 2023. Common and unique neural mechanisms of social and nonsocial conflict resolving and adaptation. Cerebral Cortex 33, 3773–3786. 10.1093/cercor/bhac306

75. Wang, R., Liu, M., Cheng, X., Wu, Y., Hildebrandt, A., Zhou, C., 2021. Segregation, integration, and balance of large-scale resting brain networks configure different cognitive abilities. Proc. Natl. Acad. Sci. U.S.A. 118, e2022288118. 10.1073/pnas.2022288118

76. Wang, S., Peterson, D.J., Gatenby, J.C., Li, W., Grabowski, T.J., Madhyastha, T.M., 2017. Evaluation of Field Map and Nonlinear Registration Methods for Correction of Susceptibility Artifacts in Diffusion MRI. Front. Neuroinform. 11. 10.3389/fninf.2017.00017

77. Wei, W., Benn, R.A., Scholz, R., Shevchenko, V., Klatzmann, U., Alberti, F., Chiou, R., Wassermann, D., Vanderwal, T., Smallwood, J., Margulies, D.S., 2024. A function-based mapping of sensory integration along the cortical hierarchy. Commun Biol 7, 1593. 10.1038/s42003-024-07224-z

78. Yan, C.-G., Craddock, R.C., Zuo, X.-N., Zang, Y.-F., Milham, M.P., 2013. Standardizing the intrinsic brain: Towards robust measurement of inter-individual variation in 1000 functional connectomes. NeuroImage 80, 246–262. 10.1016/j.neuroimage.2013.04.081

