## Supplementary Table 9 for "Normative Deviations Reveal Task-Evoked and Clinical Network Reorganization"

**Supplementary Tables and Figures**

GLM analysis

Region-wise first-level GLMs were fitted to parcellated fMRI time series (TR = 1.0 s, for more details about the parcellation see the method section of this paper). Two task regressors (NWL-task and Control-task) were constructed from condition onsets and durations and convolved with the canonical Glover hemodynamic response function. Low-frequency drifts were modeled using a cosine basis (high-pass filter: 1/128 Hz), and an intercept was included. Six rigid-body motion parameters and the global signal were added as nuisance regressors. First-level models were estimated using an AR(1) noise model.

A first-level contrast comparing conditions (task − control task) was computed for each Region. Subject-level contrast effects were subsequently entered into a second-level one-sample GLM to test whether the mean contrast effect differed from zero across subjects. Results are reported as Region-wise effect estimates, t-statistics, z-scores, and two-sided p-values.

| label | effect | t | z | p | variance |
|---|---|---|---|---|---|
| LH_SomMotB_Cent_2 | 0.056995952 | 4.609582527 | 3.730275811 | 9.56E-05 | 0.000152885 |
| aCAU-lh | 0.069981442 | 3.161434387 | 2.798162691 | 0.002569711 | 0.000490002 |
| LH_DorsAttnA_SPL_2 | 0.06024468 | 2.981167805 | 2.666044231 | 0.00383748 | 0.00040838 |
| LH_DorsAttnA_SPL_1 | 0.060083461 | 2.73165256 | 2.476943594 | 0.006625642 | 0.000483792 |
| LH_ContA_PFCl_1 | 0.048674856 | 2.687204218 | 2.442494654 | 0.007293073 | 0.000328101 |
| RH_ContC_Cingp_1 | 0.037378854 | 2.644194929 | 2.408941224 | 0.007999437 | 0.000199832 |
| LH_DefaultB_PFCv_2 | 0.030681929 | 2.493776825 | 2.289895356 | 0.011013692 | 0.000151374 |
| NAc-core-lh | 0.052577304 | 2.47734238 | 2.276728868 | 0.011401211 | 0.000450427 |
| LH_ContA_PFCl_2 | 0.076535451 | 2.476162225 | 2.275782177 | 0.011429525 | 0.00095536 |
| RH_DorsAttnA_SPL_2 | 0.050616866 | 2.42488356 | 2.234491532 | 0.012725375 | 0.000435721 |
| NAc-shell-lh | 0.041108728 | 2.37729062 | 2.195896111 | 0.01404969 | 0.000299022 |
| THA-VA-lh | 0.02653034 | 2.296201315 | 2.129534934 | 0.016605014 | 0.000133495 |
| aCAU-rh | 0.049435101 | 2.249377136 | 2.090871583 | 0.018269789 | 0.000482999 |
| LH_ContA_Temp_1 | 0.028076836 | 2.22407748 | 2.069877114 | 0.019231927 | 0.000159366 |
| LH_ContC_Cingp_1 | 0.063475791 | 2.219631899 | 2.066180497 | 0.01940572 | 0.000817814 |
| LH_ContA_IPS_3 | 0.046506165 | 1.980421467 | 1.863990904 | 0.031161491 | 0.00055145 |
| LH_ContC_pCun_1 | 0.065145601 | 1.973284483 | 1.857860792 | 0.031594393 | 0.00108991 |
| LH_SalVentAttnA_FrOper_2 | 0.03447254 | 1.952363121 | 1.839858882 | 0.032894479 | 0.000311764 |
| RH_SalVentAttnB_Ins_1 | 0.029686555 | 1.854381765 | 1.754918827 | 0.039636597 | 0.000256284 |
| LH_SalVentAttnA_FrMed_2 | 0.033831657 | 1.753436722 | 1.666339477 | 0.047822909 | 0.000372277 |

Supplementary Table 9. Region-wise GLM analysis of the NWL-task data. t= t-statistics, z = z-scores, p = two-sided p-values.
